# Reshaping the dynamics of follicle-stimulant hormone receptor models in polyunsaturated lipid bilayers. Calculation of conformational free energy landscapes of α-helical domains from all-atom MD simulations

**DOI:** 10.1101/2022.06.06.494945

**Authors:** Eduardo Jardón-Valadez, Alfredo Ulloa-Aguirre, Tobías Portillo-Bobadilla, Geiser Villavicencio-Pulido

## Abstract

G-coupled-protein receptors (GPCR) are conspicuous target molecules for novel therapeutic drugs due to their role as mediators of cellular responses. Structural biology of GPCR revealed that intracellular signaling stimulated by extracellular ligands involves subtle conformational changes of the receptor during activation. Nonetheless, transitions among intermediates evolve in an intricate and rough free energy landscape of the conformational space. Experimental evidence suggests that the membrane environment is an active modulator of the receptor dynamics; therefore, the lipid composition may facilitate conformational transitions towards productive signaling states. In this study, we setup molecular dynamics simulations to examine the conformational dynamics of the transmembrane domains, in the context of a membrane of polyunsaturated phospholipid molecules, for a homology model of the human follicle-stimulating-hormone receptor (FSHR) and the crystal structure of the Lumi intermediate of the squid rhodopsin (LSRh). The conformational dynamics of the α−helical domains of LSRh was consistent with interactions stabilizing the crystal structure, which remained well preserved in the membrane environment. In contrast, conformations in the FSHR model evolved towards stable states in the membrane environment. To assess the relevance of the conformational dynamics in the FSHR model, dihedral restraints were imposed for the helical domains on top of the force field. This strategy was implemented to reoptimize the interhelical interactions probably overlooked in the modeling process. The conformational dynamics in the helical domains was evaluated by the TM-score, contact maps, principal components analysis of Cα atoms at the helical domains, and projections of the conformational free energy on principal components. The roughness of the conformational landscape in the FSHR model without dihedral restraints, suggested that alternative interhelical conformational states were populated, whereas imposing restraints led to a dominant conformational state. Template-based models of GPCR, with reoptimized interhelical interactions using dihedral restraints, may enhance the identification of binding sites for potential therapeutic drugs.

## Introduction

G-coupled protein receptors (GPCR) encompass the largest family of membrane proteins that share a similar topology: seven-helix transmembrane (TM) domains, intra- and extracellular loops linking the TM helices, and the amino and carboxyl termini. These receptors represent highly attractive target molecules for drug discovery because of their role as mediators of an array of cellular responses triggered by a number of structurally diverse ligands including ions, photons, odorants, lipids, hormones, and neurotransmitters that vary considerably in size from small biogenic amines to peptides to large proteins [1–4]. From the outstanding structural data available in the Protein Data Bank [5], progress in the structural biology of GPCR revealed features on the conformational changes following agonist binding, intermediary stages upon receptor activation, and changes provoked by activated heterotrimeric G-proteins and other membrane-associated intracellular proteins (e.g. β-arrestins) [6]. Active conformations in crystal structures suggest similar intracellular motions in several GPCRs, including the β2-adrenergic receptor (β_2_AR), the muscarinic acetylcholine receptor (M2R), and the μ-opioid receptor (μOR), where the transmembrane (TM) helix 6 moves up to 14 Å away from the helical bundle, and TM helices 5 and 7 show a slight rotation together with an inward motion [7]. Crystal structures also revealed tight interactions involving hydrophobic pockets, H-bonds, salt bridges, and location of structural water molecules or other molecules such as detergents required for crystallization [8]. Nonetheless, finding the proper set of crystallization conditions may require up to thousands of trials, and years of intensive research [9]. Computational methods provide practical alternatives for the development of working models useful as templates for finding new agonists and antagonists, which may potentially serve as commercial drugs in pharmaceutical industry [3, 10, 11].

Molecular dynamics (MD) simulations have contributed to an atomistic description of conformational changes in membrane proteins such as the GPCRs. Analysis on trajectories at the ∼10^3^ ns time scale disclosed some features on the activation mechanisms of the β_2_AR [12], and the serotonin-2A receptor [13], including the conserved TRP switch, the opening of the ionic lock at TM helices 3 and 6, and the dynamics of the NPxxY motif present in TM helix 7. To overcome the computer time limitations to sample the conformational space, metadynamics techniques were proposed for describing the activation mechanism of GPCR [14, 15]. The emerging consensus, therefore, suggests that conformational transitions of GPCR reach 10^3^-10^6^ ns time scales [16, 17], which might be inaccessible in some cases given the system size in explicit membrane environments. At the ∼10^2^ ns time scale, MD studies on GPCR still provide valuable descriptions on fluctuations observed in the secondary structure dynamics, H-bond dynamics involving internal water molecules, side chains contacts, loop dynamics, and lipid-protein contacts [18].

Our research group has been interested in understanding the physiopathogenesis of dysfunctional gonadotropin receptors due to point mutations in the receptor protein, from the biochemical and biophysical perspectives [19–21]. Studies on the glycoprotein hormone receptors subfamily of GPCR have been addressed for the follicle-stimulating hormone receptor (FSHR), which is the prototype member of these receptors [22, 23]. Our studies include gene sequencing to identify point mutations, receptor expression in cell lines, binding to ligands and effectors, internalization and recycling of the receptor-ligand complex, and intracellular signaling [24–27]. In addition, we implemented a computational approach to determine the impact of point mutations in the interhelical region of the FSHR [28]. Interestingly, the conformational dynamics of the wild-type receptor (FSHR-WT), was sensitive to point mutations such as D408Y and D408A, according to an analysis of autocorrelation matrices of the helical domains [28]. Apparently, when replacing D408 by tyrosine or alanine, fluctuations in protein backbone atoms showed different patterns in correlated communities in a dynamical network analysis [28]. More recently, new findings in the naturally occurring I423T missense mutation located at the TM helix 2 of the FSHR (bearing the frequent polymorphic variant S680), demonstrated modestly-reduced expression at the cell surface plasma membrane and, more importantly, a significant impact of the replacement on cAMP/protein kinase A and β-arrestin-dependent ERK1/2 phosphorylation signaling [29]. According to our computational analysis, the helical integrity was well preserved in all phenotypes; nonetheless, deviations from the initial receptor coordinates were detected in some TM domains [28]. To examine in more detail the significance of conformational fluctuations in the helical domains of the FSHR-WT model, in this study we extended the simulation trajectories of the receptor to the ∼10^2^ ns time scale. In addition, we setup the crystal structure of the lumi intermediate of squid rhodopsin (LSRh) using the same membrane environment and computational protocols set for the FSHR-WT. The LSRh belongs to the rhodopsin family of GPCR, and exhibits ∼40% homology with the consensus sequence of the TM domains of the glycoprotein hormone receptors (Figure 1), corresponding to the previous intermediate of the meta I state [30].

**Fig 1.**
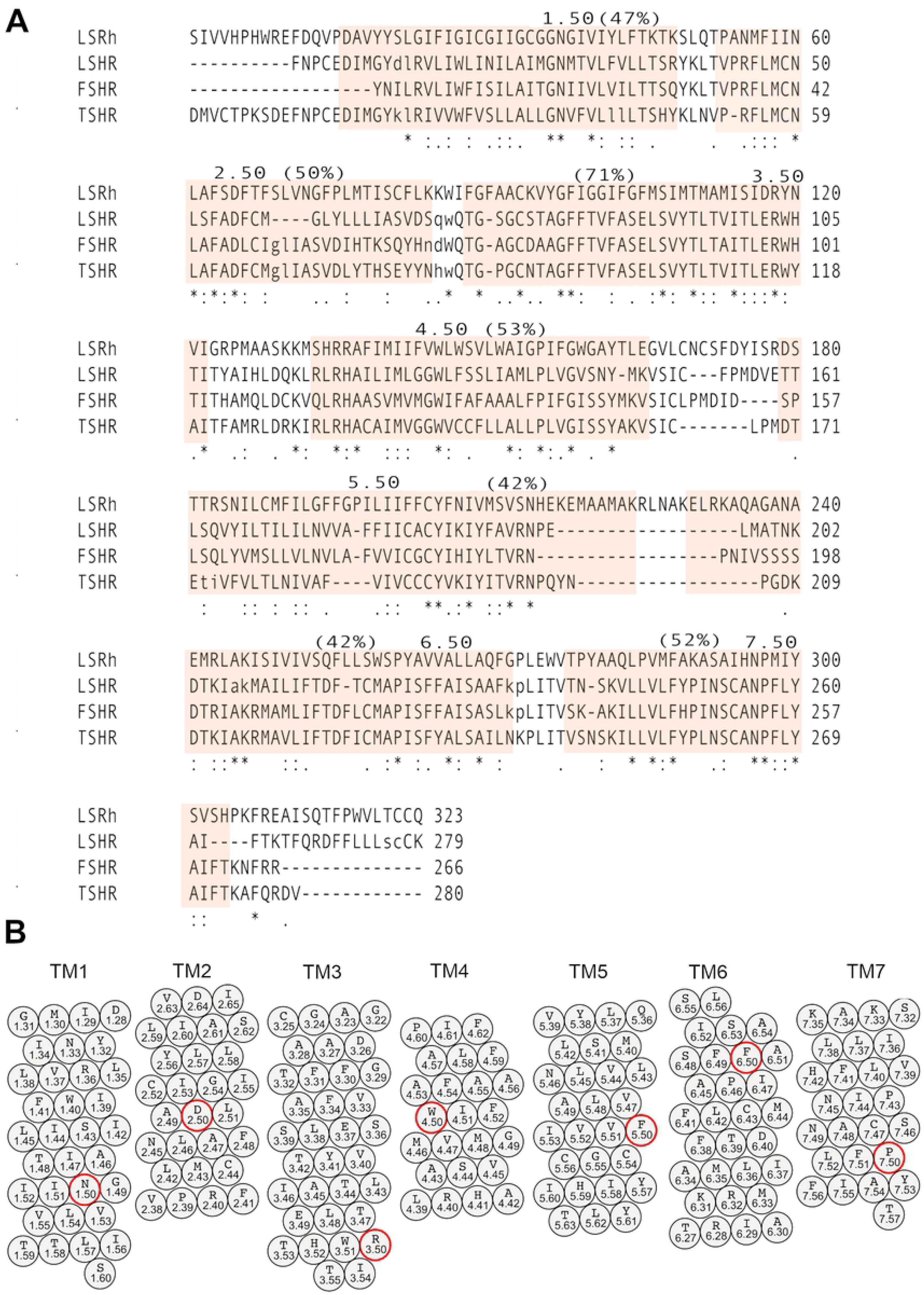
A) Sequence alignment of the squid-rhodopsin (LSRh) and the glycoprotein family: thyroid stimulant hormone receptor (TSHR), follicle stimulant hormone receptor (FSHR), and luteinizing hormone receptor (LHR). The Ballesteros and Weinstein numbering system was used, which assigns 50 to the most conserved position on TM helices 1-7 [56]. B) Schematic representation of the seven TM domains [57]. Conserved position 50 of on each TM helix is highlighted (red circle).

Alphafold2 showed accurate predictions for native protein folding [31] in the 14th Critical Assessment of protein Structure Prediction (CASP14). Nonetheless, structure-function implications in missense mutations could be beyond the applicability of the Alphafold2 workflow [32]. Thus, among other strategies to model and detect structure-function implications of point mutations, MD simulations at all-atom force-field approximation provide a physicochemical perspective to describe the significance of disruptive interactions that may impact on the native receptor structure. In this study, we setup template-based models of the FSHR-WT and the I423T mutant [29], in a lipid bilayer of polyunsaturated 1-stearoyl-2-docosahexaenoyl-sn-glycero-3-phosphocholine (SDPC) phospholipid molecules. For comparing the conformational dynamics, trajectories for the LSRh crystal structure were generated under similar conditions. We assumed that in the fluid membrane environment, interhelical interactions remain stable in the LSRh structure; in the FSHR model, interhelical interactions evolved through a rough conformational space. Previews analysis of trajectories showed that stable fluctuations could populate states compatible with the roughness of the conformational landscape of the helical domains [28]. To test the relevance of conformational dynamics in template-based models, we imposed dihedral constraints in the FSHR models on backbone atoms of the helical domains. Analysis of structural fluctuations and principal components (PC) calculation of the interhelical domains, revealed the larger extent of the conformational dynamics in the unrestrained *vs* the restrained state of the receptor. By using dihedral constrains, the FSHR model preserved the predicted helical regions in the modeling process, which was evaluated by the TM-score [33]. We conclude that our computational implementation provides a strategy to explore the structural-functional implications of missense mutations, and that it may be useful to develop new therapeutic drugs to recover receptor function in dysfunctional GPCR.

## Materials and Methods

### Preparation of simulation boxes

Simulation boxes containing the GPCR protein, SDPC lipid molecules forming the membrane, water molecules for hydration of the lipid heads, and Na^+^ or Cl^-^ ions for system neutrality were prepared similarly. Modeling of the FSHR-WT was performed as described previously [27, 28]. Briefly, the human FSHR sequence D317 to N695 was uploaded to the GPCR-I-TASSER server [34], selecting the highest scored model for subsequent refinement by MD simulation. The GPCR-I-TASSER modeling procedure includes multiple sequence alignments and experimental information stored in the GPCR database such as contact maps, as suggested by mutagenesis experiments [35]. In addition, the GPCR-I-TASSER performs *ab-initio* helix modeling, which improves the TM-score for helical domains in GPCR [34]. The initial coordinates corresponded to the highest TM-scored (0.4 ± 0.14) model calculated for the helical domains. Then, the FSHR-WT (S680) model was inserted in a pre-equilibrated lipid bilayer of SDPC molecules obtained from previous simulations [36]. Importantly, interhelical water molecules stable in the squid rhodopsin setup were preserved [37]. Disulfide bonds were defined between cysteines C338-C356 and C442-C517. Palmitoylated tails were bonded to cysteines C644 and C646 by forming tioester bonds [27]. For the mutant I423T, we used the initial receptor coordinates, and isoleucine side chain at 423 was replaced by the threonine side chain, using the mutate command in the PSFGEN script [38]. Initial box dimensions were 90 Å x 105 Å x 114 Å, respectively, for the x-, y- and z-axis. A total of 128979 atoms were included: 29231 water molecules, 248 SDPC lipid molecules, a sodium ion for charge neutrality, and the receptor FSHR-WT or mutant I423T, with 379 residues encompassing D317 to N695 [28].

The initial coordinates of the LSRh were taken from the crystal structure PDB:4WW3 chain A [39]. A pre-equilibrated membrane lipid bilayer of SDPC lipids was taken from a previews simulation of the dark state [36, 37]. The interhelical water molecules resolved in the crystal structure were preserved for the setup. Disulfide bonds between cysteines C96-C184, and C108-C186 were preserved as exhibited by the corresponding bonds in other members of the rhodopsin family of GPCR. A palmitoylated cysteine was defined at C337 and retinal was bound to K305. Initial box dimensions were 88 Å x 102 Å x 120 Å, respectively, for the x-, y- and z-axis. A total of 112118 atoms were included in the LSRh simulation box, with 23705 water molecules, 249 SDPC phospholipids, and two chloride ions for charge neutrality.

Dihedral constraints were imposed in the FSHR-WT or mutant I423T, for the helical domains, using the ssrestraints plug in of VMD [40], with options h-bonds, a dihedral force constant of 200 kcal/mol rad^2^, and including helix 1: 362 to 392; helix 2 : 397 to 432; helix 3 : 437 to 472; helix 4 : 487 to 507; helix 5: 532 to 557; helix 6: 567 to 597; and helix 7: 607 to 626 as the selected helical regions for the starting FSHR model. Helical restraints were also included for the I423T setup; therefore, restrained and unrestrained runs were executed for FSHR-WT and the I423T mutant (Table 1).

**Table 1.** Trajectory lengths for the simulation boxes of ∼120 thousand atoms containing the receptor, solvent water molecules, and SDPC lipids.

| System | Trajectory length<br>ns | Trajectory length<br>helical-<br>restraints<br>Ns | Temperature |
| --- | --- | --- | --- |
| LSRh | 145.9 |  |  |
| FSHR | 158.3 | 160.5 | 300 K |
| FSHR | 144.3 |  |  |
| I423T | 180.8 | 140.0 | 310 K |

### Force Field parameters

CHARMM36 all-atom force field parameters were used for the lipid molecules [41] and the protein atoms [42, 43], including CMAP correction [44, 45]. The retinal parameters used in the LSRh setup corresponded to those reoptimized in reference [46], which include atom charges from reference [47]. Water molecules were modeled using the TIP3P potential [48].

### Molecular Dynamics details

All simulations were performed with the NAMD 2.12 software [49]. Simulation trajectories were generated in the isothermal-isobaric ensemble (NPT) with Langevin dynamics to maintain a constant temperature, and Nosé-Hoover Langevin piston to maintain a constant pressure of 1 bar. Anisotropic cell fluctuations in the x-, y- and z-axis were allowed. Non-bonding interactions were calculated with a cutoff of 11.5 Å, and a shifting function starting at 10.0 Å. A multiple time step integration for solving the motion equations was used with one step for bonding interaction, two steps for short range nonbonding interactions, and four steps for electrostatic forces. All hydrogen atoms were fixed using the SHAKE algorithm [50]. Electrostatic interactions were evaluated using PME [51], with a 4^th^ order interpolation on a grid of ∼1 Å in the *x-*, *y-* and *z-*directions, and a tolerance of 10^-6^ for the direct evaluation of the real part of the Ewald sum. The unrestrained trajectories were generated for the FSHR-WT at 300K as well as at the physiological temperature of 310 K (Table 1). Trajectory of the FSHR-WT at 300 K correspond to *in-vitro* experimental conditions [29]. Trajectories using dihedral constrains were generated for FSHR-WT at 300K, and I423T at 310 K. Trajectory of the I423T mutant at 310 K using dihedral constrains was expected to recover interhelical interactions at physiological temperature upon disruption of side chain contacts. The LSRh trajectory was generated without helical restraints at 300K; LSRh system was considered as a comparable reference system given that it shows ∼40% of sequence homology with the glycoprotein hormone subfamily of GPCR (Fig 1). The set of simulations were prepared to explore the conformational space of the seven-helix domain in the FSHR-WT, and the novel mutant I423T [29].

### Trajectory analysis

Conformational changes were calculated by the root mean square deviation (RMSD) for the Cα atoms of the helical domains in FSHR-WT and I423T mutant: Y362 to Y392 (helix 1), P397 to Y432 (helix 2), Q437 to T472 (helix 3), A487 to G507 (helix 4), M532 to R557 (helix 5), D567 to L597 (helix 6), and A607 to Y626 (helix 7). In LSRh: D31 to T62 (helix 1), P68 to L98 (helix 2), A106 to G138 (helix 3), H149 to P170 (helix 4), D194 to R240 (helix 5), L246 to P288 (helix 6), and P294 to V317 (helix 7). For each TM helix, contact maps were calculated for side chain heavy atoms using a cutoff of 3.5 Å. Hence, TM interactions were detected as function of simulation time whenever side chain heavy atoms were within the cutoff distance.

To assess deviations from the initial helical structure of the transmembrane domains, we calculated the TM-score defined as:

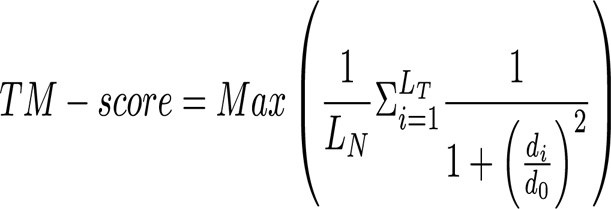

where L_N_ is the length of the reference structure, L_T_ is the length of the aligned residues, d_i_ is the distance between the *i-th* pair of aligned residues, and d_0_ is a scale to normalize the match difference. ‘Max’ denotes the maximum value after optimal spatial superposition. The TM-score takes values within 0 and 1, with TM-score > 0.5 for well-preserved helical domains [33].

The conformational landscapes of LSRh, FSHR-WT, and I423T were obtained by a principal component analysis of the Cα carbons (N) using the GROMACS 2019.4 suite of programs [52, 53]. By diagonalization of covariance matrices, the resultant matrix transformation contained eigenvectors in 3 x N columns as well as the eigenvalues corresponding to the mean square fluctuation in direction of the principal modes, and ordered from the largest to the lowest contribution to the total fluctuation [54]. As a measure of the total fluctuation, the root mean square fluctuation (RMSF) was calculated from the square root of the sum of eigenvalues divided by the number of Cα atoms (N). The simulations trajectories were divided in blocks of 2^9^, 2^10^, …, 2^16^ ps in order to calculate averages and standard deviations as a function of the block size [55]. Visualization of the trajectory projected over the first six PC was generated using the VMD software [40]. Free energy landscapes were calculated from projections of the principal modes, PC1 to PC6. Our computational approach, including the system set-up, generation of MD trajectories, and analysis of fluctuations allowed us to apply a strategy to evaluate both the conformational dynamics of a membrane receptor relaxed from a crystal structure, such as the LSRh, or a homology model for the FSHR-WT, and the impact of the I423T mutation located at the TM helix 2 of the helical bundle

## Results

By comparing the conformational dynamics of LSRh and a homology model of the FSHR-WT and I423T mutant, we analyzed the dynamics of the Cα atoms at the helical domains (Fig 1). Such strategy was implemented to assess the conformational fluctuations as a means to account for the biological relevance of helical motion in hormone receptor models *vs* a photon-activated x-ray structure (Fig 2). From a previous study, the integrity of the helical domains of the FSHR-WT was evaluated according to calculations of the secondary structure, number of residues forming each TM helix and root mean squared deviations (RMSD) of each TM helix, RMSF, and other parameters. We found that helical domains were well preserved [28]. Nonetheless, using the TM-score we detected values below the 0.5 helicity threshold for both, the FSHR-WT and I423T mutant, suggesting a disruption of the TM folding of these FSHRs (Fig 3). Interestingly, the TM-score in LSRh showed values close to one for TM helices 1, 2, 5, and 6, whereas TM helices 3, 4, and 7 exhibited significant variability, yet above the 0.5 threshold. TM-score values for TM helices in LSRh suggest that the secondary structure was well preserved. As the initial LSRh structure was obtained from x-ray diffraction at 2.8 Å resolution [39], helical interactions were compatible with the tight packing of the crystal structure, and remained stable in the fluid environment of the membrane. Importantly, TM-score calculations for LSRh were performed without any restrained force on top of the force field.

**Fig 2.**
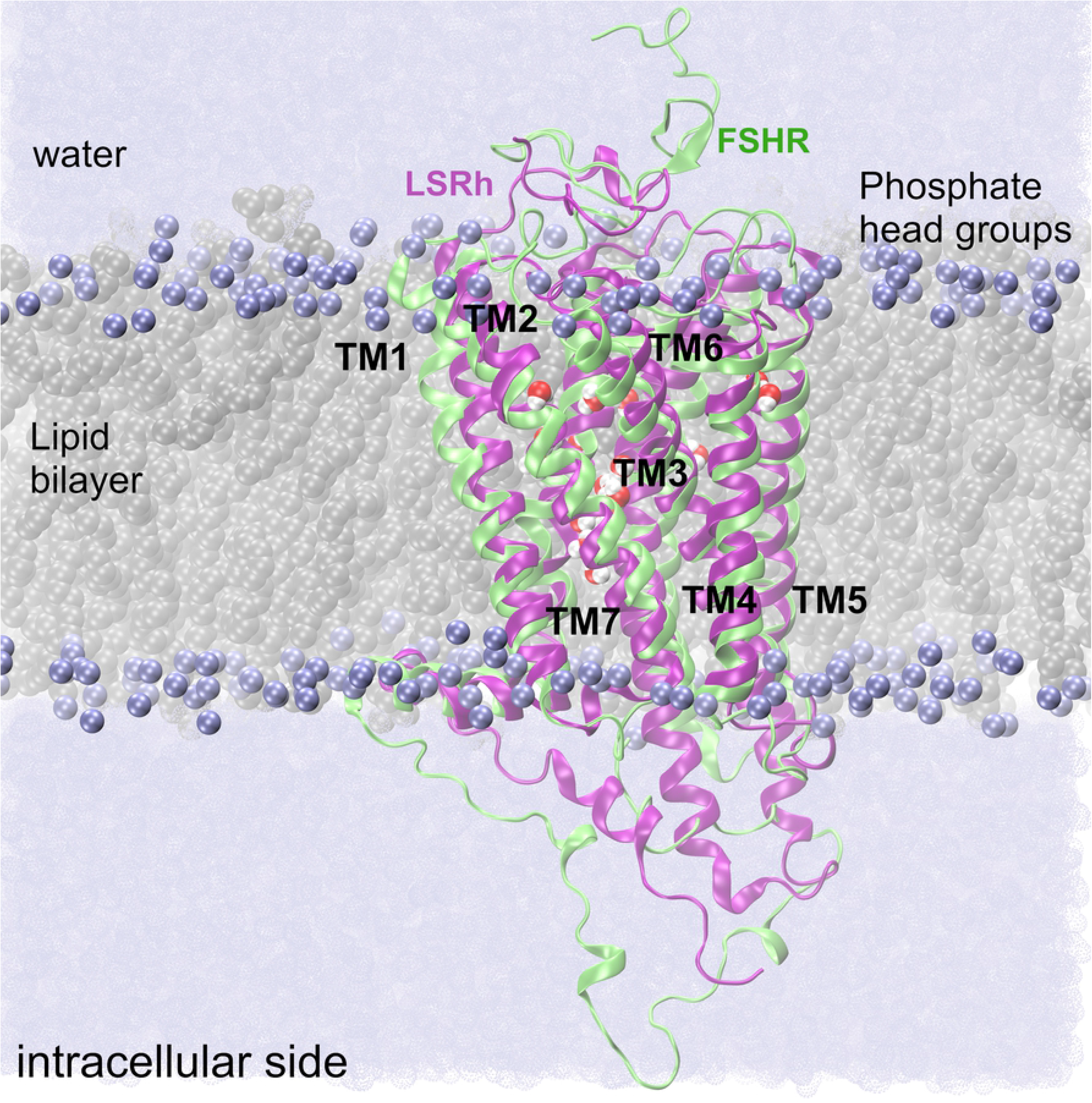
Membrane receptors in the lipid bilayer environment at 300 K. FSHR-WT and LSRh are depicted, respectively, as green and magenta ribbons. SDPC lipid molecules are depicted as spheres in the background, with carbon atoms in gray, and phosphate atoms as ice-blue. The solvent water medium is depicted as a continuum in ice blue. The structural alignment of the receptors shows consistency in the location of the helical domains in the hydrophobic region of the membrane. The structural alignment between these receptors show some differences, for example, the extension of intracellular domains of TM5 and TM6, the tilt angle of TM1, and kink angle of TM7.

**Fig 3.**
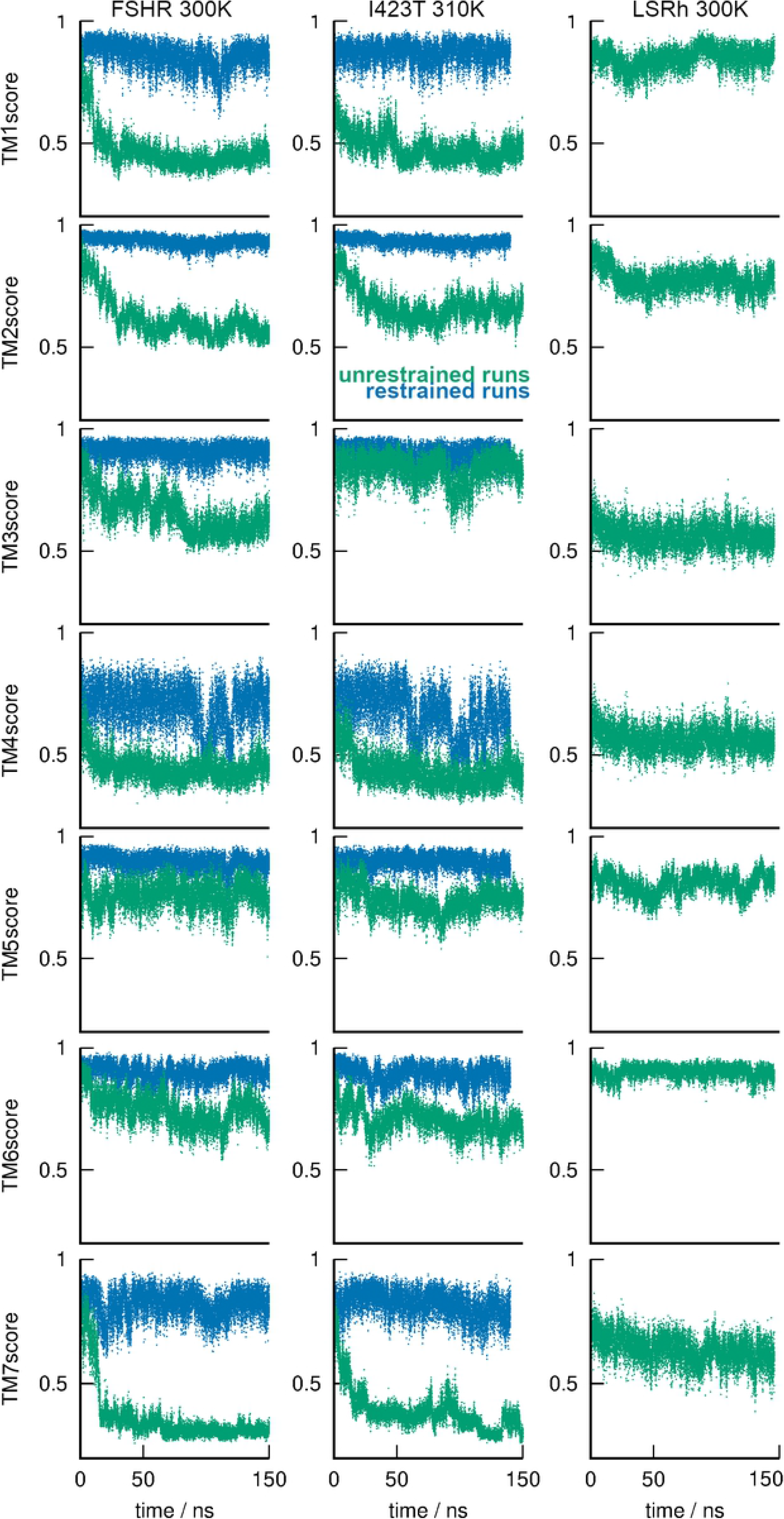
TM-score of the TM domains of the FSHR-WT and I423T models, respectively at 300 and 310 K, and the LSRh at 300K. Unrestrained runs correspond to dash lines in green. Blue dash lines represent the TM-score for the restrained runs. TM-score showed well reserved helicity in the restrained runs.

In FSHR-WT and I423T, the TM-score values displayed significant differences depending on the simulation conditions. On the one hand, TM-score values for TM helices 1, 4, and 7 were lower than 0.5 in the unrestrained runs (Table 2); for TM helices 2, 3, 5, and 6 TM-score the corresponding values fluctuated above the 0.5 helicity threshold (Table 2). Interestingly, TM-score values of helix 5 were similar to those calculated for the LSRh run (Fig 3). On the other hand, in the restrained runs all TM-score values were stable and close to one (Table 2). TM helices 1, 4, 6 and 7 showed similar trends to those calculated for the LSRh (Fig 3). For TM helix 5, our analysis suggested that it might not necessary to use helical restraints at all, since the TM-score values fluctuated as those in LSRh (Table 2 and Fig 3). In TM helix 2, the TM-score was larger in the receptor models than the values calculated for LSRh (Table 2 and Fig 3). Regarding the magnitude of the force constant of the helical restraints, TM-score variability was well reproduced as that observed in LSRh, except for TM helix 2; for example, FAHR-WT helix 1, 4, 6, and 7 showed fluctuations within magnitudes similar to those in LSRh (Table 2 and Fig 3). In TM helix 2, a weaker force constant might improve the comparison against helix 2 of LSRh. Thus, TM-score was a parameter sensitive to distortions of helicity in the TM domains in the FSHR-WT and I423T mutant. Finally, the response of TM-score due to the increase in temperature from 300 K and 310K in FSHR-WT showed similar values within error bars, except for TM helix 7 of FSHT-WT, whose helicity was better preserved at 300K (S1 Figure in S1_info, Table 2, and Fig 3). Similar trends in the TM-score of I423T at 310K were detected, except for the higher helicity of TM helix 3 relative to that in FSHR-WT at 300K (Table 2, and Fig 3). The overall results showed that helicity measured by the TM-score was higher in LSRh relative to those in FSHR-WT and I423T without helical restraints, which suggested that the intermolecular interactions stabilizing the helical domains in LSRh were well represented by the force field; hence, helicity disruption in the receptor models such as the FSHR-WT and I423T were largely related to suboptimal interhelical interactions.

**Table 2.** TM-score values for each TM helix in restrained and unrestrained runs, and at 300 K and 310 K. Values larger than 0.5 represent well preserved helicity.

| System | TM-score |  |  |  |  |  |  | Temp |
| --- | --- | --- | --- | --- | --- | --- | --- | --- |
|  | TM1 | TM2 | TM3 | TM4 | TM5 | TM6 | TM7 |  |
| LSRh | 0.87±0.02 | 0.76±0.03 | 0.80±0.03 | 0.60±0.05 | 0.79±0.04 | 0.86±0.02 | 0.71±0.04 | 300 K |
| FSHR | 0.46±0.03 | 0.54±0.03 | 0.64±0.04 | 0.46±0.04 | 0.73±0.05 | 0.71±0.05 | 0.31±0.02 |  |
| FSHR+ restraints | 0.86±0.04 | 0.93±0.02 | 0.91±0.03 | 0.72±0.07 | 0.89±0.03 | 0.91±0.03 | 0.82±0.04 |  |
| FSHR | 0.51±0.03 | 0.59±0.05 | 0.70±0.05 | 0.41±0.04 | 0.77±0.04 | 0.73±0.05 | 0.53±0.03 | 310 K |
| I423T | 0.49±0.04 | 0.67±0.03 | 0.81±0.06 | 0.40±0.04 | 0.73±0.04 | 0.69±0.05 | 0.32±0.03 |  |
| I423T+ restraints | 0.87±0.04 | 0.93±0.01 | 0.90±0.03 | 0.66±0.08 | 0.88±0.03 | 0.89±0.04 | 0.79±0.05 |  |

Dihedral restraints had an impact on the conformational changes according to RMSD calculations of the TM domains. Fig 4 shows time series for the RMSD for all runs. After 20 ns of trajectory RMSD fluctuated at an average of 1.2 ±0.10 Å in LSRh, and 0.83 ± 0.11 Å and 0.86 ±0.12 Å, respectively, for the restrained runs of FSHR-WT and I423T. For the unrestrained runs, after 50 ns of trajectory, RMSD values reached an average 1.75±0.19 Å and 1.99 ± 0.12, respectively, for FSHR-WT and I423T at 310 K; for FSHR-WT at 300 K, RMSD reached 2.42±0.15 Å. The larger RMSD averages in unrestrained runs were expected because restrained forces attenuated displacements of TM domains. To measure the positional displacements from the starting structure, RMSD calculations for each TM helix were also performed. In the restrained runs, RMSD averages were calculated from 0.29 to 0.51 Å for the FSHR-WT helices, and 0.31 to 0.53 for the I423T mutant (S2 and S3 Figures in S1_info). In LSRh, RMSD averages from 0.6 to 1.2 Å suggested slight positional displacements of the TM helices in the fluid environment of the membrane (S4 Fig in S1_info). Without restrained forces, TM helices showed stable average configurations in both, FSHR-WT and I423T at 310 K: RMSD from 0.6 to 1.2 Å for helices 3, 4, 5, and 6; and RMSD 1.3 to 2.0 Å for helices 1, 2, and 7. Larger RMSD values for individual TM helices correlated with decreasing helicity, for example, in helices 1 and 7 (Fig 3).

**Fig 4.**
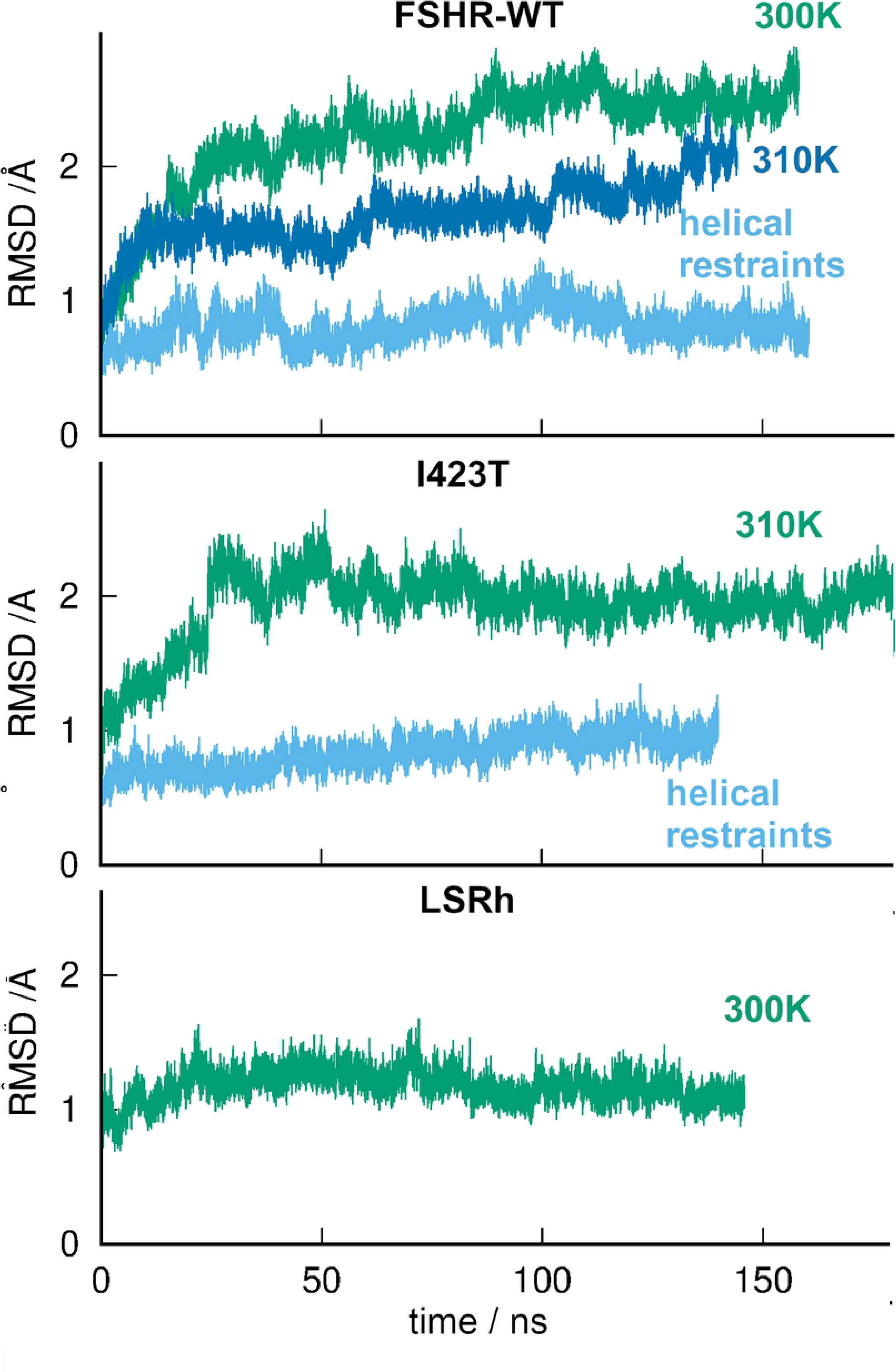
Root mean square deviations (RMSD) of the helical domains in the FSHR-WT (top plot) and I423T mutant (mid plot), with and without helical restraints, at 300 K or 310K. RMSD for the LSRh helical domains at 300K (bottom plot). Using helical restraints modulate the positional displacements.

Interactions stabilizing the 3D structure in the FSHR-WT and I423T models were evaluated by measuring contact maps for each TM-helix. Importantly, TM helices work as molecular switches since interhelical interactions may lead to conformational changes [58]. Fig 5 shows the results for contact maps in the FSHR-WT for the restrained and unrestrained runs. Consistently detected contacts were exhibited by the TM helical domains: TM1 made permanent contacts with TM2 and TM7; TM2, with TM3, TM4, and TM7; TM3, with TM4, TM5, and TM7; TM4, with TM5; TM5, with TM6; and TM6, with TM7. In TM2 of I423T mutant, contacts with TM7 involved V612, H615, L616, S619, Y626, T630, and F633 (Fig 6), with a difference in the restrained *vs* the unrestrained run consisting in fewer partner exchanges and interhelical contacts detected every 3 or 4 residues apart. Contacts between TM7 and side chains of TM6 were absent in the restrained run, for example with side chains of M576, T580, and K598 (Fig 6). TM3 showed significant contacts with TM6 in the unrestrained run, involving L577, D581, C584, and M585, and I588 (Fig 6). For the light receptor LSRh, Fig 7 shows contacts among the TM helices: TM1 was in permanent contact with TM2, and TM7; TM2, with TM3, and TM7, and the intracellular half of TM4; TM3, with TM4, TM5, and TM6; TM5, with TM6; and TM6, with TM7. In summary, contacts detected in the LSRh were consistent with contacts identified in the FSHR-WT and I423T models, although slight differences were detected in restrained *vs* the unrestrained runs. No disruption of interhelical contacts due to a temperature increase were detected, as replica of the FSHR-WT at 310 K showed similar interhelical contacts as trajectory at 300 K (S5 Fig in S1_info).

**Fig 5.**
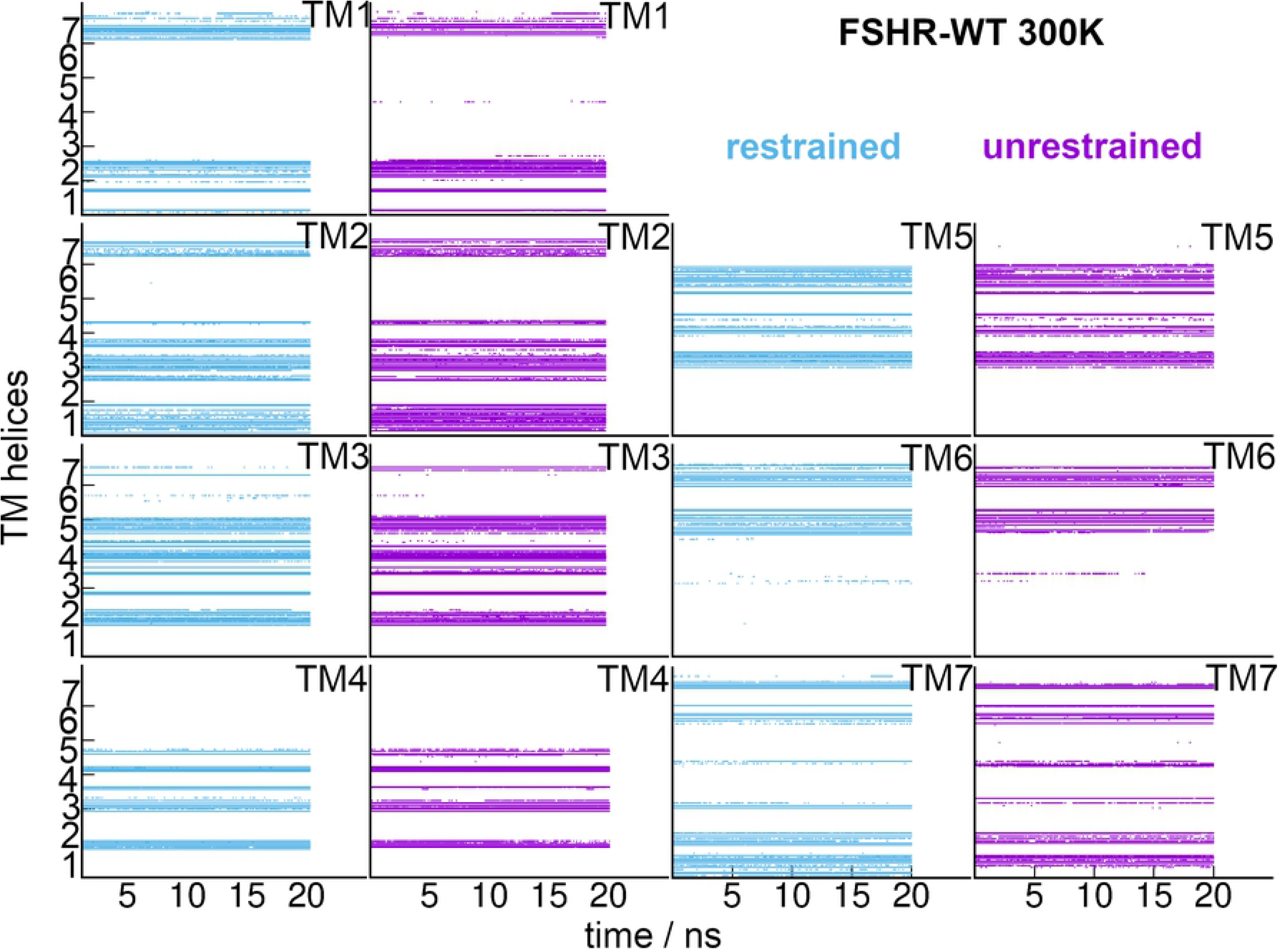
Contact maps as function of simulation time for the TM-helices in the FSHR-WT at 300 K. Data points were depicted whenever residues of TM-X were at 3.5 Å of residues of TM-Y. In the vertical axis the TM helices were identified from 1 to 7, for each plot of TM1-7. Analysis performed for the last 20 ns of trajectory.

**Fig 6.**
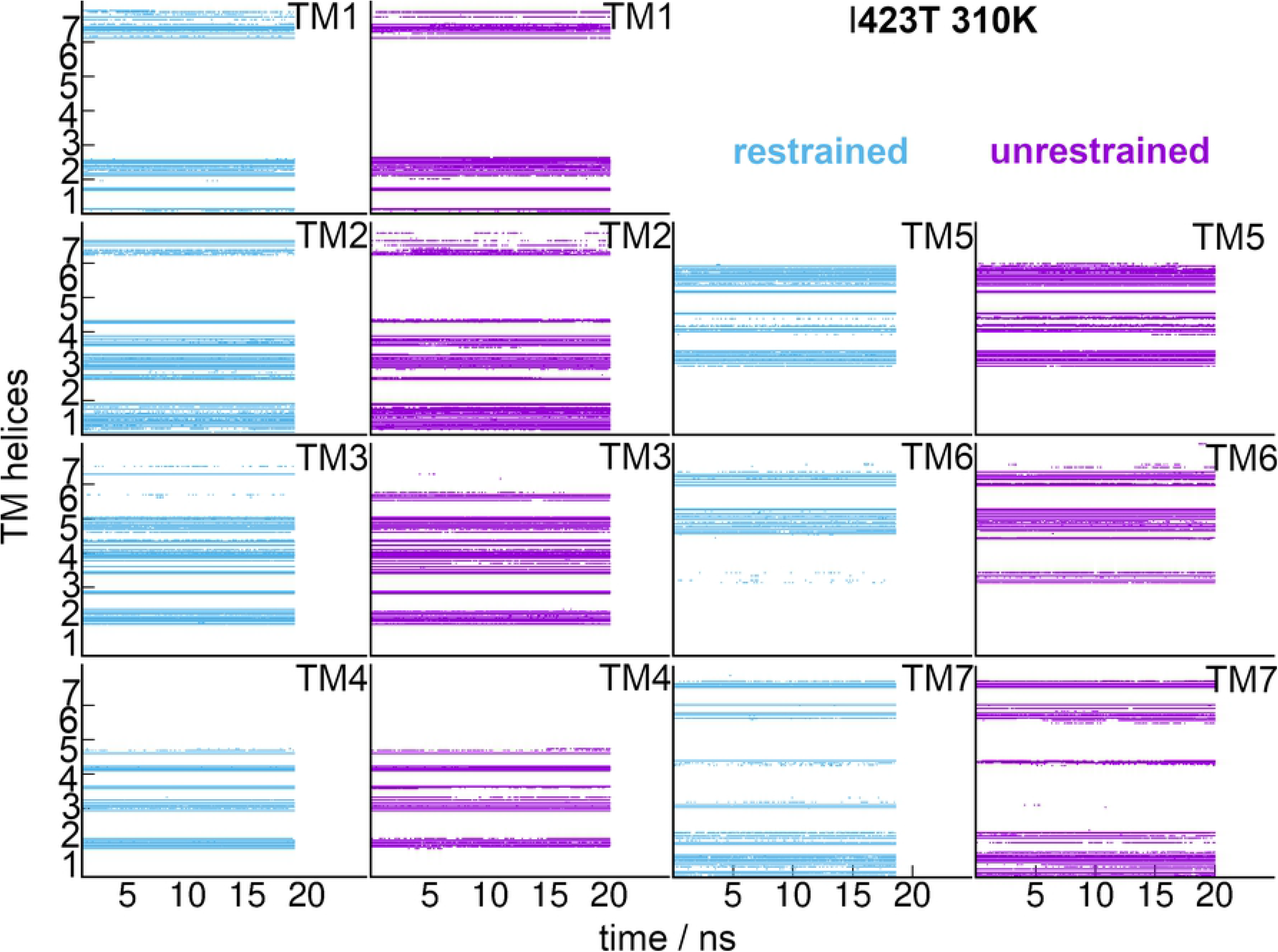
Contact maps as function of simulation time for the TM-helices in the I423K at 310 K. Data points were depicted whenever residues of TM-X were at 3.5 Å of residues of TM-Y. In the vertical axis the TM helices were identified from 1 to 7, for each plot of TM1-7. Analysis performed for the last 20 ns of trajectory.

**Fig 7.**
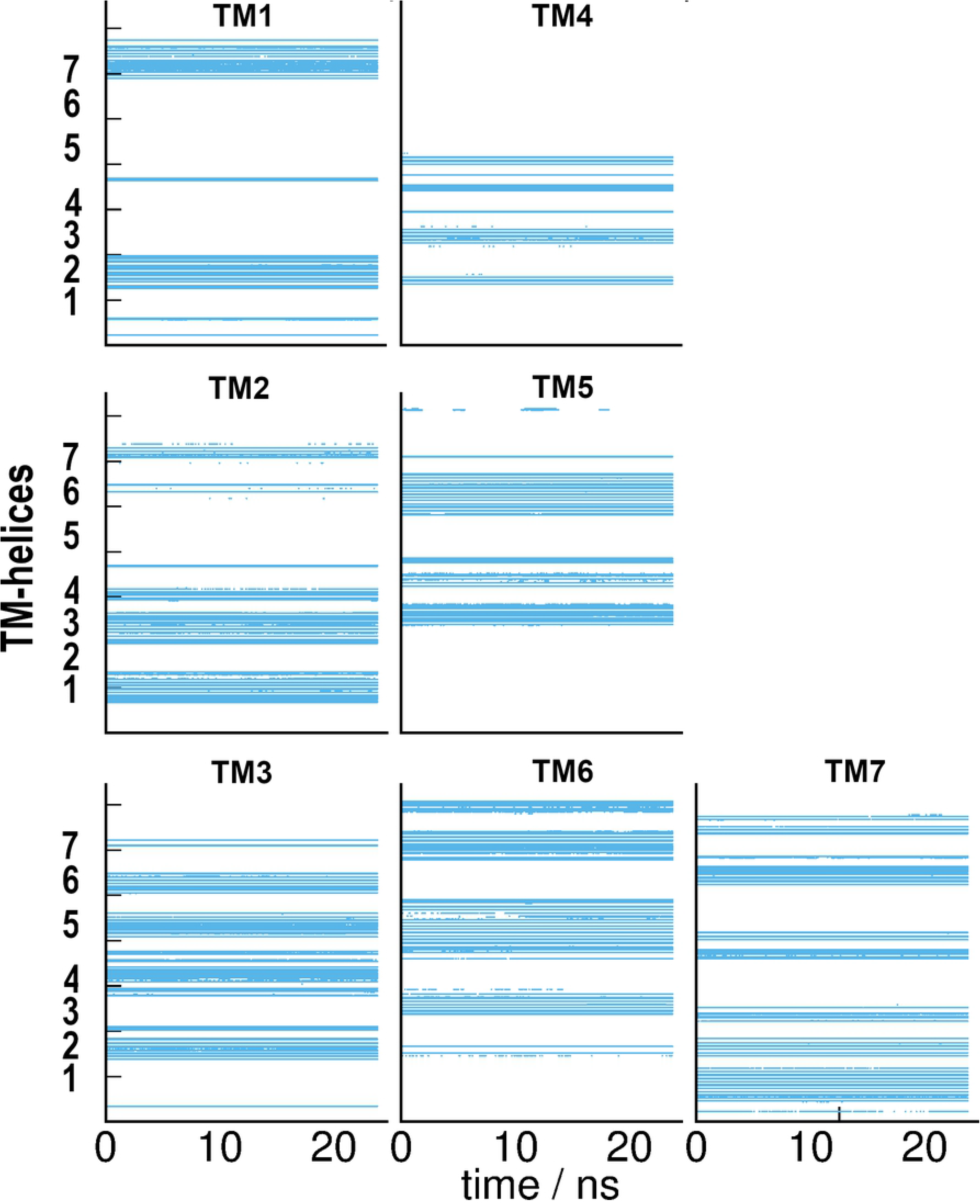
Contact maps as function of simulation time for the TM1-TM7 of LSRh. Data points depicted whenever residues of TM-X were at 3.5 Å of residues of TM-Y. In the vertical axis the TM helices are identified as from 1 to 7, for each plot of TM1-7. Analysis performed for the last 20 ns of trajectory. TM1 was in contact mainly with TM2 and TM7; TM2 made contact with TM3, and TM7; TM3 with TM4-6; TM5 with TM4 and TM6; TM6 with TM7.

Because of the helical restraints imposed on FSHR-WT and I423T mutant, conformational states would differ from those sampled in the unrestrained runs. Even though the unrestrained runs preserved the overall structure of the receptor [28], in present study we examined more deeply the relevance of conformational fluctuations of receptor models using the Cα atoms at the helical domains only. Fig 8 shows distributions for PC1-6 of the FSHR-WT, and I423T mutant obtained from PCA calculations; the normalized histograms for restrained *vs* unrestrained runs were overlapped for comparisons. Profiles of the restrained runs showed unimodal distributions, except in PC2 of FSRL-WT and PC1 of I423T, which showed distributions with two and three maxima, respectively. Unrestrained runs showed wider distributions than those obtained for the restrained trajectories, which suggests that the receptor evolved to states not accessible when helical restraints were imposed (Fig 8). Fig 9 shows the conformational change due to PC1, which corresponds to the motion that includes the largest contribution to the total fluctuation. In LSRh and the restrained FSHR-WT, helical motions showed small amplitudes in all TM helices, with consistent displacements in TM helices 1-4, as well as the extracellular half of TM5 and TM6 (Fig 9). The main difference between LSRh and restrained FSHR-WT was observed in the intracellular side of TM of helices 5 and 6: in LSRh, TM5 moved toward TM6, while TM6 moved inwards whereas in FSHR-WT, both TM5 and TM6 displaced outwards. The PC1 collective motion in restrained I423T showed small amplitudes in all helical domains, except in TM6 which moved outwards. In the unrestrained runs of FSHR-WT at 300 K and 310 K, displacements were consistent in amplitudes and directions with those observed in the restrained run, although the upper half of TM6 moved with larger amplitude at 310 K (Fig 9). Despite the slight displacements in the 3D space, the transformation of the covariance matrix performed in PCA allowed to identify the axis (eigenvectors) that account for the variability of the Cα coordinates. In the rotated coordinates, the PC disclosed the collective low frequency modes of motion for the helical domains. Simulation studies on resolved structures of GPCR performed at the micro- and millisecond time scales provided some clues on the activation mechanism, such as the dynamics of the conserved TRP switch and the NPxxY motif, and the opening of the ionic lock at TM3 and TM6 [12, 13]. Such long simulation times scales were required to overcome configurational free energy barriers [59, 60]. Using a subset of coordinates such as the Cα atoms of the helical domains, at the 10^2^ ns time scale, the LSRh trajectory was expected to relax into a dominant conformational intermediary, which was evident from the calculation of unimodal distributions in PC1-6 (Fig 10). Therefore, from the profiles in the distributions of the PC1-6 in the FSHR-WT and I423T models, and the LSRh, we noticed that the unrestrained runs showed significant differences in the dynamics of the helical domains, probably related to suboptimal interhelical interactions.

**Fig 8.**
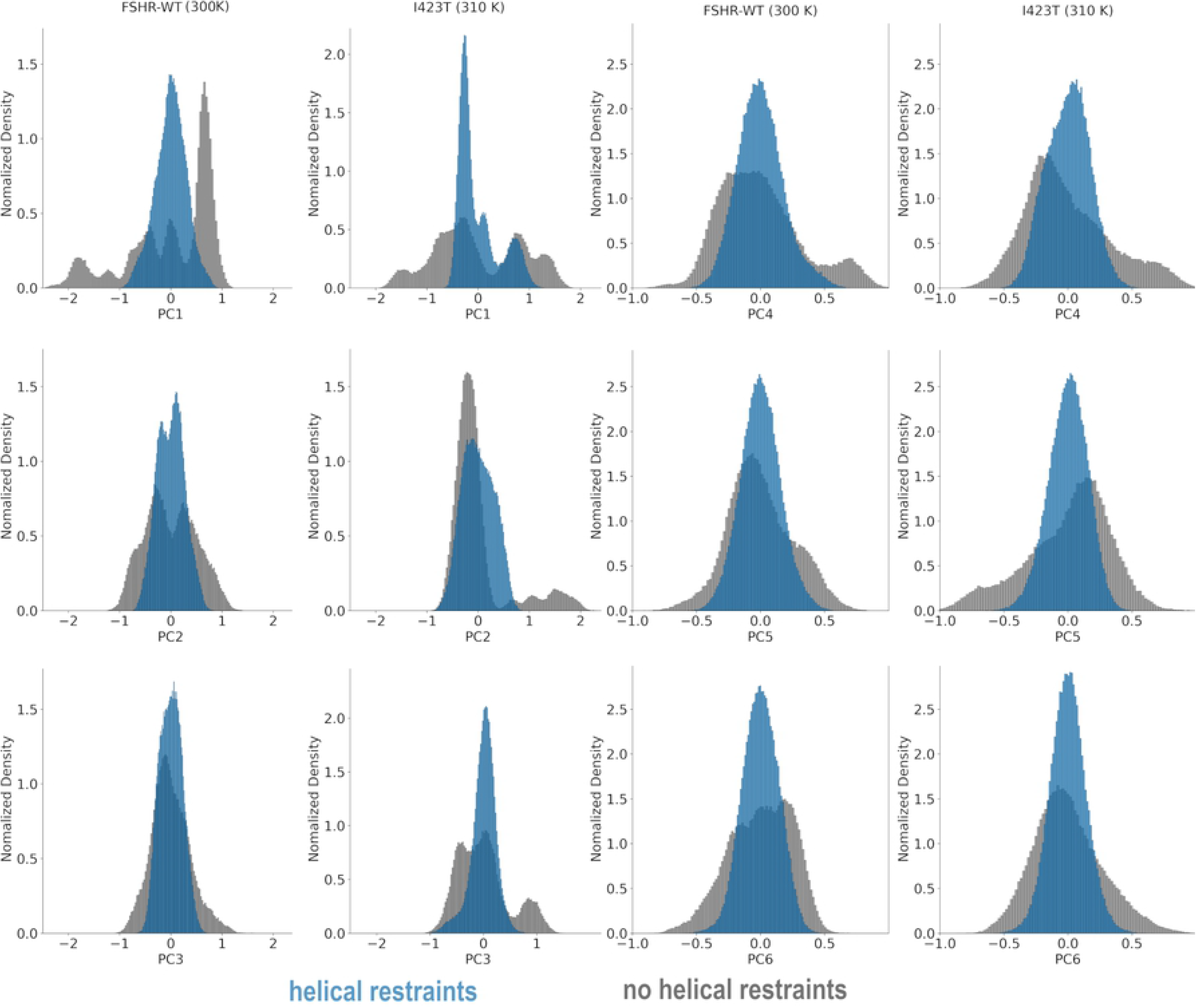
Normalized density distributions for vectors PCA1-6 (nm) for the FSHR-WT, and the I423T mutant. Distributions for the restrained and unrestrained runs are shown. Unimodal distributions were found in the restrained runs, except for TM2 of FSHR-WT and TM1 of I423T. PC distributions unveil the population of conformational states sampled in the simulation trajectories.

**Fig 9.**
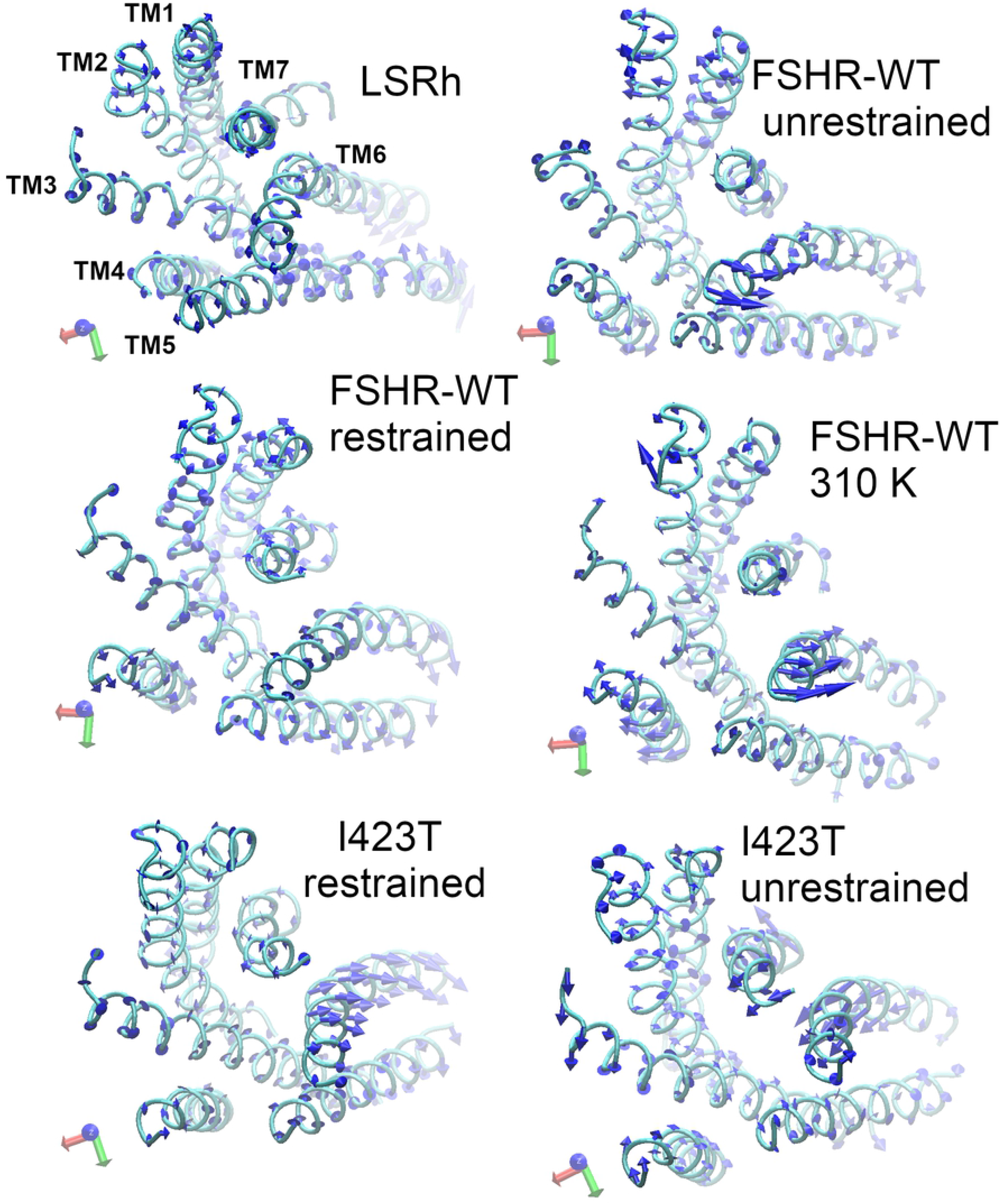
Top view of helical domains, TM1-TM6, for receptors LSRh, FSHR-WT and I423T. For a reference coordinates x, y, and z are shown, respectively, as red, green, and blue arrows; all views share the same relative position of TM helices as in panel A. A. Helical domains of the LSRh, helices TM1 to TM6, are depicted as cyan ribbons. Blue arrows depicted on the Cα atoms represent the direction of the motion associated to the first principal component whose sizes correspond to the magnitude of the displacement. B. Helical domains of the FSHR-WT receptor model, at 300K. C. Helical domains of the FSHR-WT including helical restraints, at 300K. D. Helical domains of the FSHR-WT receptor model, at 310K. E. Helical domains of the I423T receptor model including helical restraints, at 310K. F. Helical domains of the I423T receptor model at 310K.

**Fig 10.**
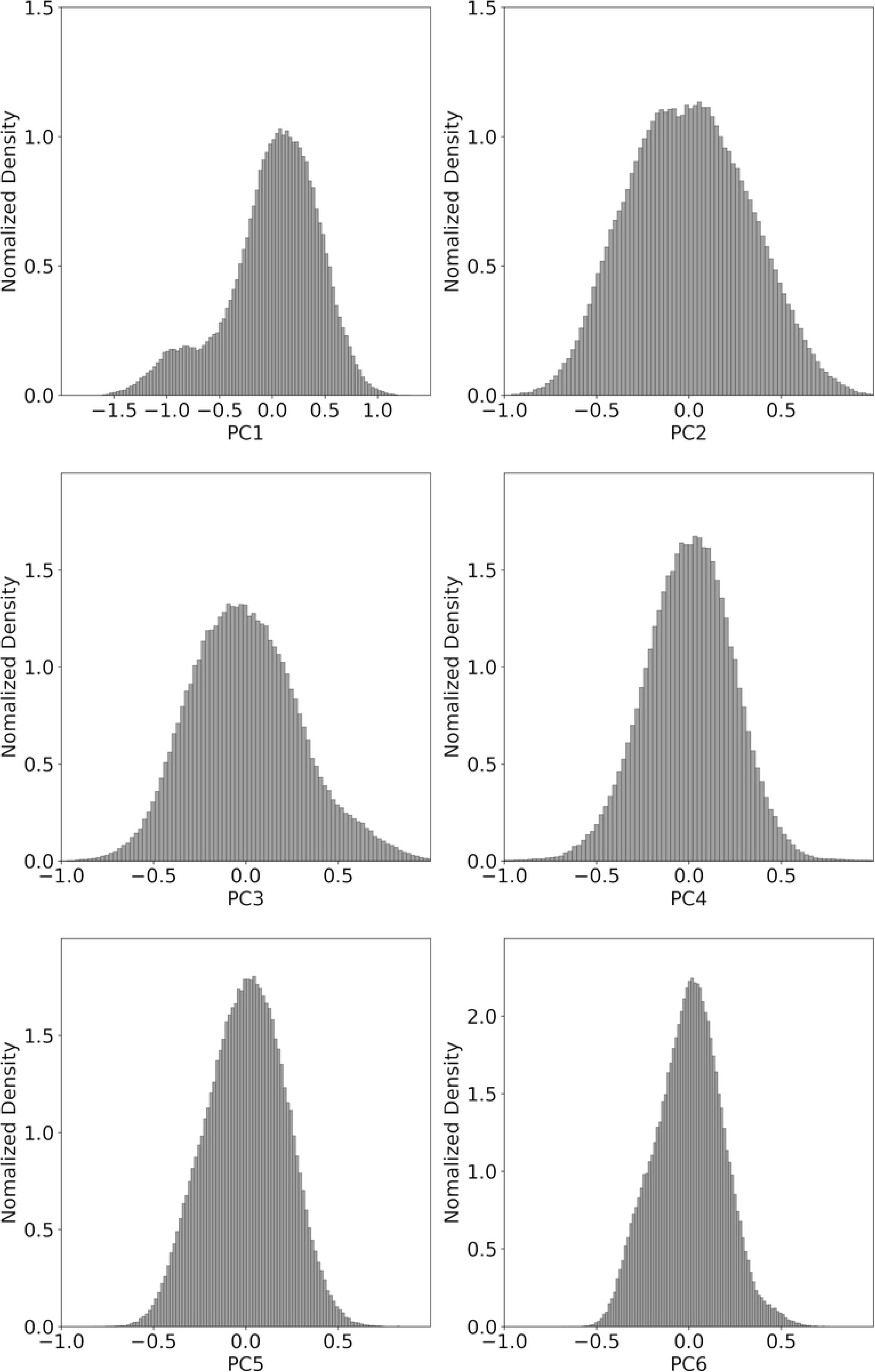
Normalized density distributions for vectors PCA1-6 (nm) for the LSRh. PC distributions unveil the population of conformational states sampled in the simulation trajectories. Unimodal distributions of the LSRh suggests that fluctuations were detected for one main conformational state.

Further, from the calculation of the total fluctuation (RMSF) of Cα atoms in blocks of 2^9^ to 2^16^ time frames (1 ps each), we determined the block size for the convergence of the RMSF (S6 Figure in S1_info). In LSRh, total fluctuation reached constant value of 4 pm for 2^16^ time frames (65.5 ns). In contrast, for the unrestrained runs, the total RMSF increased with the block size. When helical restraints were imposed, the total RMSF in both FSHR-WT and the I423T showed a similar behavior as the LSRh, reaching a 3 pm plateau for a block size of 2^16^ time frames. In summary, the PCA1-6 of unrestrained runs showed broad distributions, whereas the restrained runs as well as LSRh showed mostly unimodal distributions in the PC1-6 principal modes.

By choosing specific atoms of the protein structure we implemented PCA to determine the relevance of positional fluctuations expected for atoms at the helical region in a light activated receptor and in hormone receptor models. Including the overall 3D structure of a membrane protein to detect conformational changes may require long time scales to sample a complex configurational space. Nonetheless, PCA analysis on subdomains (e.g. binding pocket residues, backbone atoms, etc.) may provide characteristics of the dynamics on a local scale, even at the ∼10^2^ ns time scale [18]. Interesting, projections of the free energy landscape corresponding to the PCA of a reduced set of 3D coordinates, such as the Cα atoms of the helical domains, provide an energetic criterium for the transitions among different conformations generated by principal components. Figures 11 and 12 show the free energy projections of PC1-2, PC3-4, and PC5-6 for FSHR-WT and I423T models, respectively. For comparisons, the free energy projections for the LSRh were included in both figures. In Fig 8 we calculated multipeak probability density distributions for the unrestrained runs mainly for PC1-3, which produced several local free energy minima spread in the PC1-PC2 and PC3-PC4 projections (Figs 11 and 12). In the unimodal distributions of Fig 8, corresponding to restrained runs, free energy minima in all projections were localized in a central region of the plots (Figs 11 and 12). For the LSRh trajectory, projections of free energy showed well defined minima in PC3-4 and PC5-6, whereas few local minima were detected in PC1-2; transitions among configurations due to PC1-2 could be expected since no helical restrains were imposed on the LSRh structure. Therefore, from projections of the free energy on PC, we could determine the extent of the conformational dynamics of FSRH-WT and I423T models in a fluid membrane environment, relative to the dynamics of the LSRh. By using dihedral restraints in the FSRH-WT and I423T models, the modulation of helical dynamics allowed to preserve interhelical contacts as well as the integrity of the helicity of the TM domains, which are known to work as structural switches on the activation process of GPCR. Our results suggest that including helical restraints could improve the quality of a working model for membrane receptors, such as the FSHR-WT, without any additional cost of computer time in comparison to the generation of unrestrained trajectories.

**Fig 11.**
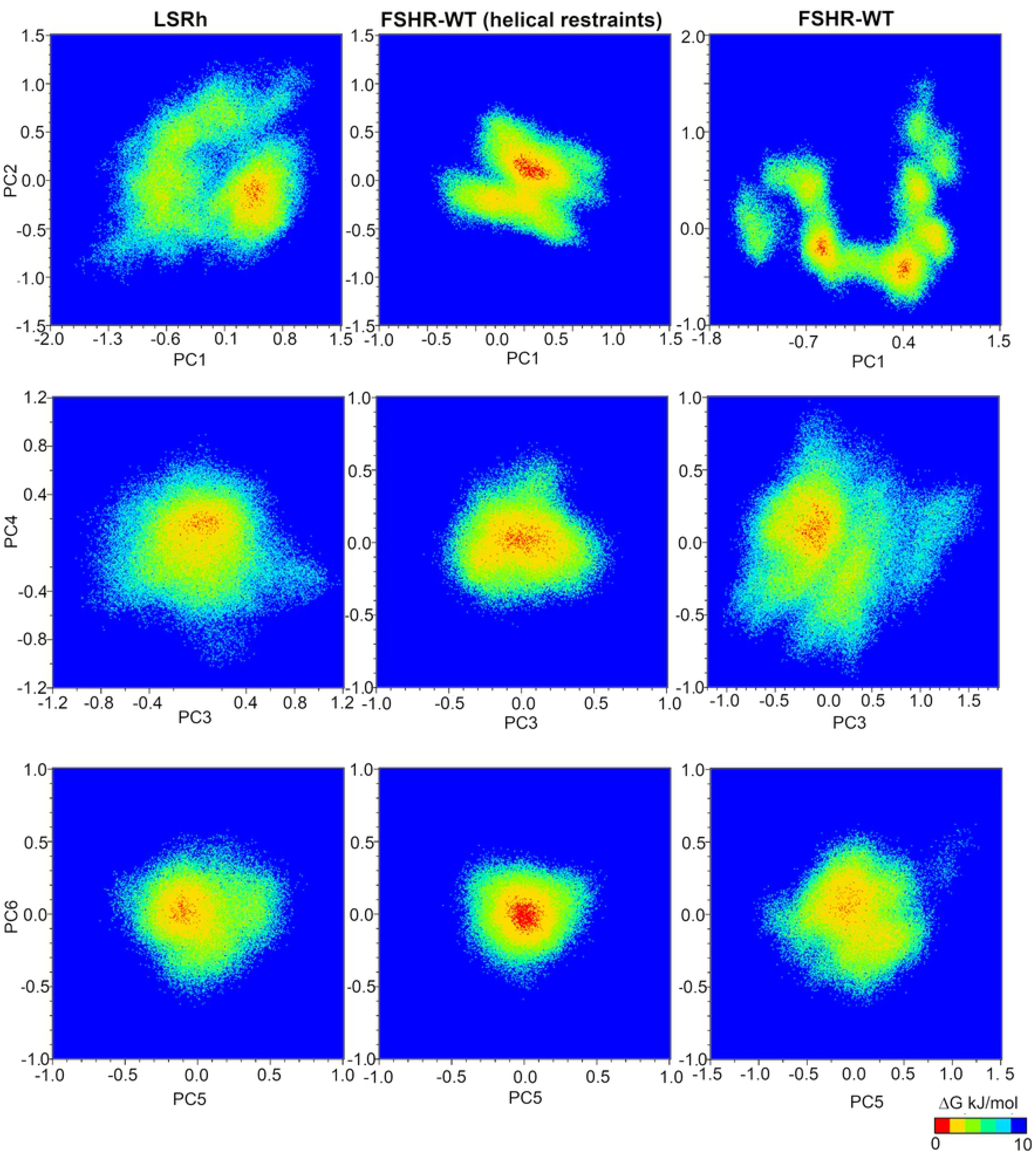
Projection of the free energy landscape on principal components, vectors 1 to 6 (in nm), for LSRh and FSHR-WT at 300 K. Red dots cluster each other in the free energy minimum. In the unrestrained run of FSHR-WT, the projection of PC1 and PC2 disclose a broad distribution of local minima that suggests transitions among several conformational states.

**Fig 12.**
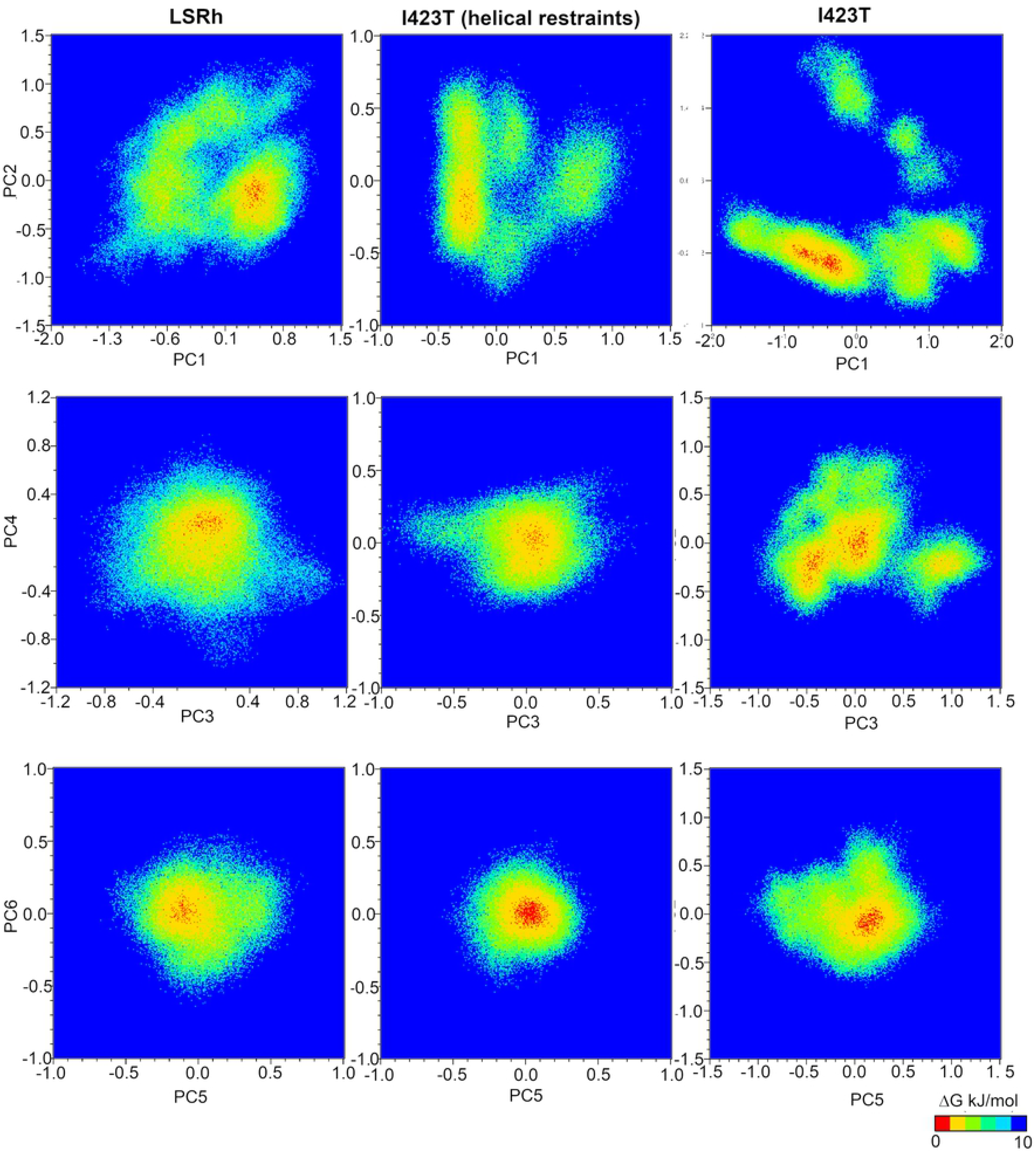
Projection of the free energy landscape on principal components, vectors 1 to 6 (in nm), for LSRh at 300 K, and I423T at 310 K. Red dots cluster each other in the free energy minimum. In the unrestrained run of I423T, the projection of PC1 and PC2 disclose a broad distribution of local minima that suggests transitions among several conformational states.

## Discussion

Understanding signal transduction mediated by the FSHR, a prototype member of the glycoprotein hormones subfamily of GPCR, relies on the description of the conformational free energy landscape in the membrane environment. In the process of signal internalization, the intracellular receptor domains engage G proteins, or β-arrestins [6] through conformational changes that expose particular intracellular sequences involved in the activation of these pivotal, universal signal transducers [7, 14, 16]. In this study, we used the covariance matrix to unveil the conformational dynamics of helical domains in simulation trajectories at the ∼10^2^ ns time scale for both the FSHR-WT and the I423T mutant at 300 and 310 K, respectively (Table 1). Further, to evaluate the significance of conformational fluctuations in template-based models, the dynamics in a homologous receptor (Figs 1 and 2) such as the LSRh were also analyzed. Relaxation of the initial LSRh crystal structure in a fluid membrane environment, evolved to states compatible with intermediates in early stages of activation. Therefore, we chose the LSRh structure as a reference system because it is an early intermediate of the activation of visual rhodopsin in invertebrates [39], with the retinal chromophore in all-*trans* configuration. Hence, it was expected that fluctuations of the TM domains, helices 1 to 7, populate configurational states in preparation for the conformational changes forming the binding pocket for engaging intracellular transducers. Finding a dominant state in a relaxed crystal structure or in a reoptimized model could suggest common features, assuming that TM interactions emerge from: I) Sequence similarity, whereby side chain interactions could be found in conserved motifs; II) Interhelical interactions that became permanent in the simulation not necessarily present in the initial model; and III) Interhelical interactions in the crystal structure that remain stable in the fluid membrane environment. By using dihedral restraints in the FSRH-WT and I423T models, the modulation of helical dynamics allowed to preserve interhelical contacts as well as the integrity of the helicity of the TM domains, which are known to work as structural switches on the activation process of GPCR. From total RMSF calculated for the Cα atoms, we found that 2^16^ time frames were sufficient to include most of the variability in the restrained runs, while in the unrestrained runs the total RMSF was not stable at the ∼10^2^ time scale. Our results suggest that including helical restraints could improve the quality of a working model for membrane receptors, such as the FSHR-WT, without any additional cost of computer time in comparison to the generation of unrestrained trajectories.

## Conclusions

In this study we explored the conformational dynamics of the LSRh relaxed in a membrane bilayer as well as a working model of the FSHR-WT and its I423T mutant. Since the homology modeling protocol employed was based on the GPCR-Tasser approach —which takes into account the TM-score as a parameter to evaluate the quality of a GPCR model— we tested the behavior of such parameter as a function of the simulation time for a set of MD trajectories at 300 K and 310 K, including helical restraints in the FSHR-WT and I423T models. We found that using helical restraints improves the TM-score with values close to one, without compromising the computer time required to generate simulation trajectories at ∼10^2^ ns time scale. The advantages of using helical restraints in working models included the predominant conformational state shown by the dynamics of the receptor, resembling the behavior also detected for the LSRh structure and the re-optimization of the interactions among the TM helices when the homology modeling misses relevant contacts in the helical bundle. Although the overall structure of the receptor is well preserved in the unrestrained runs and the first principal mode presented similar trends in the collective motion of the TM helices, we suggest that helical restraints may be part of the modeling refinement of a GPCR homology model.

**Eduardo Jardón-Valadez:** Conceptualization, investigation, methodology, writing - review & editing. **Alfredo Ulloa-Aguirr**e Conceptualization, investigation, writing - review & editing. **Tobías Portillo-Bobadilla**: Writing - review & editing. **Geiser Villavicencio-Pulido:** Writing - review & editing.

## Acknowledgments

Authors acknowledge the access and support from supercomputing facilities: cluster YOLTLA at UAM-Iztapalapa, and cluster Axolotl at UAM-Lerma.

## Supporting information

S1_info. Data describing the dynamics of the 3D structure of the membrane protein receptors: FSHR-WT, I423T and LSRh.

**S1 Fig.** Time series for the TM-score of FSHR-WT

**S2 Fig.** Root mean square displacement (RMSD) for FSHR-WT

**S3 Fig.** Root mean square displacement (RMSD) for I423T

**S4 Fig.** Root mean square displacement (RMSD) for LSRh

**S5 Fig.** Contact maps as function of simulation time for the TM-helices in the FSHR-WT

**S6 Fig**. Root mean square fluctuations of Ca atoms calculated as function of block size for trajectories

## Notes

### Competing Interest Statement

The authors have declared no competing interest.

## References

1. Ernst OP, Lodowski DT, Elstner M, Hegemann P, Brown LS, Kandori H. Microbial and animal rhodopsins: structures, functions, and molecular mechanisms. Chemical reviews. 2014;114(1):126–63.

2. Millar RP, Newton CL. The Year In G Protein-Coupled Receptor Research. Molecular Endocrinology. 2010;24:261–74.

3. Hauser AS, Chavali S, Masuho I, Jahn LJ, Martemyanov KA, Gloriam DE, et al. Pharmacogenomics of GPCR drug targets. Cell. 2018;172(1-2):41–54.

4. Salon JA, Lodowski DT, Palczewski K. The Significance of G Protein-Coupled Receptor Crystallography for Drug Discovery. Pharmacol Rev. 2011;63:901–37.

5. Berman HM, Westbrook J, Feng Z, Gilliland G, Bhat TN, Weissig H, et al. The Protein Data Bank. Nucleic Acids Res. 2000;28(1):235–42.

6. Hilger D, Masureel M, Kobilka BK. Structure and dynamics of GPCR signaling complexes. Nature structural & molecular biology. 2018;25(1):4–12.

7. Latorraca NR, Venkatakrishnan AJ, Dror RO. GPCR dynamics: structures in motion. Chemical reviews. 2017;117(1):139–55.

8. Murakami M, Kouyama T. Crystal structure of squid rhodopsin. Nature. 2008;453:363–8.

9. Li X, Dang S, Yan C, Gong X, Wang J, Shi Y. Structure of a presenilin family intramembrane aspartate protease. Nature. 2013;493(7430):56.

10. Koehler Leman J, Ulmschneider MB, Gray JJ. Computational modeling of membrane proteins. Proteins: Structure, Function, and Bioinformatics. 2015;83(1):1–24.

11. Almeida JG, Preto AJ, Koukos PI, Bonvin AMJJ, Moreira I. Membrane proteins structures: A review on computational modeling tools. Biochimica et Biophysica Acta-Biomembranes. 2017;1859:2021–39.

12. Dror RO, Arlow DH, Maragakis P, Mildorf TJ, Pan AC, Xu H, et al. Activation mechanism of the β_2_-adrenergic receptor. Proceedings of the National Academy of Sciences of the United States of America. 2011;108:18684–9.

13. Shan J, Khelashvili G, Mondal S, Mehler EL, Weinstein H. Ligand-dependent conformations and dynamics of the serotonin 5-HT 2A receptor determine its activation and membrane-driven oligomerization properties. PLoS Comput Biol. 2012;8(4):e1002473.

14. Harpole TJ, Delemotte L. Conformational landscapes of membrane proteins delineated by enhanced sampling molecular dynamics simulations. Biochimica Et Biophysica Acta (BBA)-Biomembranes. 2018;1860(4):909–26.

15. Provasi D, Filizola M. Putative active states of a prototypic g-protein-coupled receptor from biased molecular dynamics. Biophysical Journal. 2010;98(10):2347–55.

16. Grossfield A. Recent progress in the study of G protein-coupled receptors with molecular dynamics computer simulations. Biochimica et Biophysica Acta (BBA)-Biomembranes. 2011;1808(7):1868–78.

17. Ribeiro JML, Filizola M. Allostery in G protein-coupled receptors investigated by molecular dynamics simulations. Current opinion in structural biology. 2019;55:121–8.

18. Bahar I, Lezon TR, Bakan A, Shrivastava IH. Normal mode analysis of biomolecular structures: functional mechanisms of membrane proteins. Chemical reviews. 2010;110(3):1463–97.

19. Maya-Núñez G, Janovick JA, Aguilar-Rojasa A, Jardón-Valadeza E, Leaños-Miranda A, Zariñana T, et al. Biochemical mechanism of pathogenesis of human gonadotropin-releasing hormone receptor mutants Thr104Ile and Tyr108Cys associated with familial hypogonadotropic hypogonadism. Molecularand Cellular Endocrinology. 2011;337:16–23.

20. Ulloa-Aguirre A, Zariñán T, Jardón-Valadez E. Misfolded G Protein-Coupled Receptors and Endocrine Disease. Molecular Mechanisms and Therapeutic Prospects. International Journal of Molecular Sciences. 2021;22(22):12329.

21. Jardón-Valadez E, Ulloa-Aguirre A, Piñeiro A. Modeling and Molecular Dynamics Simulation of the Human Gonadotropin-Releasing-Hormone Receptor in a Lipid Bilayer. J Phys Chem B. 2008;112:10704–13.

22. Jiang X, Dias JA, He X. Structural biology of glycoprotein hormones and their receptors: Insights to signaling. Molecular and Cellular Endocrinology. 2014;382:424–51.

23. Ulloa-Aguirre A, Zariñán T, Jardón-Valadez E, Gutiérrez-Sagal R, Dias JA. Structure-Function Relationships of the Follicle-Stimulating Hormone Receptor. Frontiers in Endocrinology. 2018;9:707. doi: 10.3389/fendo.2018.00707.

24. Garcia-Jimenez G, Zarinan T, Rodriguez-Valentin R, Mejia-Dominguez NR, Gutierrez-Sagal R, Hernandez-Montes G, et al. Frequency of the T307A, N680S, and -29G>A single-nucleotide polymorphisms in the follicle-stimulating hormone receptor in Mexican subjects of Hispanic ancestry. Reprod Biol Endocrinol. 2018;16(1):100. Epub 2018/10/21. doi: 10.1186/s12958-018-0420-4. PubMed PMID: 30340493.

25. Ulloa-Aguirre A, Zariñán T. The Follitropin Receptor: Matching Structure and Function. Mol Pharmacol. 2016;90:596–608.

26. Ulloa-Aguirre A, Reiter E, Crepieux P. FSH Receptor Signaling: Complexity of Interactions and Signal Diversity. Endocrinology. 2018;159(8):3020–35. Epub 2018/07/10. doi: 10.1210/en.2018-00452. PubMed PMID: 29982321.

27. Melo-Nava B, Casas-González P, Pérez-Solís MA, Castillo-Badillo J, Maravillas-Montero JL, Jardón-Valadez E, et al. Role of Cysteine Residues in the Carboxyl-Terminus of the Follicle Stimulating Hormone Receptor in Intracellular Traffic and Postendocytic Processing. Frontiers in Cell and Developmental Biology. 2016;4:1–13.

28. Jardón-Valadez E, Castillo-Guajardo D, Martínez-Luis I, Gutiérrez-Sagal Rn, Zariñán T, Ulloa-Aguirre A. Molecular dynamics simulation of the follicle-stimulating hormone receptor. Understanding the conformational dynamics of receptor variants at positions N680 and D408 from in silico analysis. Plos One. 2018;13(11):e0207526.

29. Zariñán T, Mayorga J, Jardón-Valadez E, Gutiérrez-Sagal R, Maravillas-Montero JL, Mejía-Domínguez NR, et al. A novel mutation in the FSH receptor (I423T) affecting receptor activation and leading to primary ovarian failure. The Journal of Clinical Endocrinology & Metabolism. 2021;106(2):e534–e50.

30. Menon ST, Han M, Sakmar TP. Rhodopsin: structural basis of molecular physiology. Physiological reviews. 2001;81(4):1659–88.

31. Jumper J, Evans R, Pritzel A, Green T, Figurnov M, Ronneberger O, et al. Highly accurate protein structure prediction with AlphaFold. Nature. 2021;596(7873):583–9.

32. Buel GR, Walters KJ. Can AlphaFold2 predict the impact of missense mutations on structure? Nature Structural & Molecular Biology. 2022;29(1):1–2.

33. Xu J, Zhang Y. How significant is a protein structure similarity with TM-score= 0.5? Bioinformatics. 2010;26(7):889–95.

34. Zhang J, Yang J, Jang R, Zhang Y. GPCR-I-TASSER: A Hybrid Approach to G Protein-Coupled Receptor Structure Modeling and the Application to the Human Genome. Structure. 2015;23:1538–49.

35. Zhang J, Zhang Y. GPCRRD: G protein-coupled receptor spatial restraint database for 3D structure modeling and function annotation. Bioinformatics. 2010;26(23): 3004–5. doi: 10.1093/bioinformatics/btq563.

36. Járdon-Valadez E, Bondar A-N, Tobias DJ. Coupling of Retinal, Protein, and Water Dynamics in Squid Rhodopsin. Biophysical Journal. 2010;99:2200–7.

37. Járdon-Valadez E, Bondar A-N, Tobias DJ. Dynamics of the internal water molecules in squid rhodopsin. Biophysical Journal. 2009;96:2572–6.

38. Gullingsrud J, Saam J, Phillips J. psfgen User’s Guide version 1.62012 2013. Available from: http://www.ks.uiuc.edu/Research/vmd/plugins/psfgen/ug.pdf.

39. Murakami M, Kouyama T. Crystal Structure of the Lumi Intermediate of Squid Rhodospin Plos One. 2015;10:e0126970.

40. Humphrey W, Dalke W, Schulten K. VMD: Visual molecular dynamics. JMolGraphics. 1996;14(1):33–8. PubMed PMID: 21602.

41. Lee S, Tran A, Allsopp M, Lim JB, Hénin J, Klauda JK. CHARMM36 United Atom Chain Model for Lipids and Surfactants. The Journal of Physical Chemitry B. 2014;118:547–56.

42. MacKerell AD, Jr., Bashford D, Bellott M, Dunbrack RL, Jr., Evanseck JD, Field MJ, et al. All-atom empirical potential for molecular modeling and dynamics studies of proteins. JPhysChemB. 1998;102(18):3586–616. PubMed PMID: 1.

43. MacKerell AD, Jr., Feig M, Brooks CL, II. Extending the treatment of backbone energetics in protein force fields: Limitations of gas-phase quantum mechanics in reproducing conformational distributions in molecular dynamics simulations. JComputChem. 2004;25:1400–15.

44. Best RB, Zhu X, Shim J, Lopes PEM, Mittal J, Feig M, et al. Optimization of the additive CHARMM all-atom protein force field targeting improved sampling of the backbone phi, psi and side-chain chi1 and chi2 dihedral angles,. Journal of Chemical Theory and Computation. 2012;8:3257–73,.

45. MacKerell AD, Feig M, Brooks CL. Improved Treatment of the Protein Backbone in Empirical Force Fields. Journal of the American Chemical Society. 2004;126(3):698–9. doi: 10.1021/ja036959e.

46. Gruia A, Bondar A-N, Smith JC, Fischer S. Mechanism of a Molecular Valve in the Halorhodopsin Chloride Pump. Structure. 2005;13:617–27.

47. Nina M, Roux B, Smith JC. Functional interactions in bacteriorhodopsin: A theoretical analysis of retinal hydrogen bonding with water. Biophysical Journal. 1995;68:25–39.

48. Jorgensen WL, Chandrasekhar J, Madura JD, Impey RW, Klein ML. Comparison of simple potential functions for simulating liquid water. JChemPhys. 1983;79(2):926–35. PubMed PMID: 21603.

49. Phillips JC, Braun B, Wang W, Gumbart J, Tajkhorshid E, Villa E, et al. Scalable molecular dynamics with NAMD. Journal of Computational Chemistry. 2005;26:1781–802.

50. Miyamoto S, Kollman P. An analytical version of the SHAKE and RATTLE algorithm for rigid water models. Journal of Computational Chemistry. 1992;13:952–62.

51. Essmann U, Perera L, Berkowitz ML, Darden T, Lee H, Pedersen LG. A smooth particle mesh Ewald method. JChemPhys. 1995;103:8577–93. PubMed PMID: 17108.

52. GROMACS 2019.4 [Internet]. 2019 [cited 2019]. Available from: https://doi.org/10.5281/zenodo.3460415.

53. Abraham MJ, Murtola T, Schulz R, Páll S, Smith JC, Hess B, et al. GROMACS: High performance molecular simulations through multi-level parallelism from laptops to supercomputers. SoftwareX. 2015;1:19–25.

54. Hess B. Convergence of samplig in protein simulations. Phys Rev E. 2001;65:031910–10.

55. Flyvbjerg H, Petersen HG. Error estimates on averages of correlated data. JChemPhys. 1989;91:461–6. PubMed PMID: 20532.

56. Ballesteros JA, Weinstein H. Integrated methods for the construction of three-dimensional models and computational probing of structure-function relations in G protein-coupled receptors. In: Sealfon SC, editor. Receptor Molecular Biology. Methods in Neurosciences: Academic Press; 1995. p. 366–428.

57. Worth CL, Kreuchwig F, Tiemann JKS, Kreuchwig A, Ritschel M, Kleinau G, et al. GPCR-SSFE 2.0—a fragment-based molecular modeling web tool for Class A G-protein coupled receptors. Nucleic acids research. 2017;45(W1):W408–W15.

58. Hauser AS, Kooistra AJ, Munk C, Heydenreich FM, Veprintsev DB, Bouvier M, et al. GPCR activation mechanisms across classes and macro/microscales. Nature structural & molecular biology. 2021;28(11):879–88.

59. Dawaliby R, Trubbia C, Delporte C, Masureel M, Van Antwerpen P, Kobilka BK, et al. Allosteric regulation of G protein–coupled receptor activity by phospholipids. Nature chemical biology. 2016;12(1):35–9.

60. Hu X, Provasi D, Ramsey S, Filizola M. Mechanism of μ-opioid receptor-magnesium interaction and positive allosteric modulation. Biophysical journal. 2020;118(4):909–21.

